# Tumor Phylogeny Topology Inference via Deep Learning

**DOI:** 10.1101/2020.02.07.938852

**Authors:** Erfan Sadeqi Azer, Mohammad Haghir Ebrahimabadi, Salem Malikić, Roni Khardon, S. Cenk Sahinalp

## Abstract

Principled computational approaches for tumor phylogeny reconstruction via single-cell sequencing typically aim to build the most likely perfect phylogeny tree from the noisy genotype matrix - which represents genotype calls of single-cells. This problem is NP-hard, and as a result, existing approaches aim to solve relatively small instances of it through combinatorial optimization techniques or Bayesian inference. As expected, even when the goal is to infer basic topological features of the tumor phylogeny - rather than reconstructing the topology entirely, these approaches could be prohibitively slow. In this paper, we introduce fast deep-learning solutions to the problems of inferring whether the most likely tree has a linear (chain) or branching topology and whether a perfect phylogeny is feasible from a given genotype matrix. We also present a reinforcement learning approach for reconstructing the most likely tumor phylogeny. This preliminary work demonstrates that data-driven approaches can reconstruct key features of tumor evolution.

## 1 Introduction

Cancer is an evolutionary disease characterized by progressive accumulation of somatic mutations in tumor cells. Deciphering the evolutionary history of a given tumor represents an important problem in studies of cancer and can help us in better understanding of several clinically important aspects of the tumor, including progression, metastatic spread, the existence of divergent subclones evolving on different branches of the tumor phylogenetic tree and many others.

Due to the importance of the problem, there have been rapid developments in the design of principled computational methods for tumor phylogeny inference. Many of these methods use bulk sequencing data where DNA from millions of cancerous and normal cells are sequenced together. Tree inference from this type of data is typically based on the use of cancer cell fraction of detected variants - in particular, single-nucleotide variants (Strino et al. 2013, Deshwar et al. 2015, El-Kebir et al. 2015, Malikic et al. 2015, Popic et al. 2015, Donmez et al. 2017, El-Kebir et al. 2016, Satas & Raphael 2017, Myers et al. 2019, Husić et al. 2019), but also copy number (e.g., (Zaccaria et al. 2017)) and structural variants (e.g., (Eaton et al. 2018, Ricketts et al. 2019)). While being cost-effective, the low resolution of bulk sequencing data is a limiting factor in tumor evolution modeling. In particular, bulk sequencing data from a single tumor sample typically admits a linear topology as an optimal solution under common tree-scoring models (Donmez et al. 2017, Gerstung et al. 2020). However, inferring whether the underlying tumor includes divergent subclones, which evolve through distinct branches of the tumor phylogeny, represents an important step towards better understanding tumor progression and improving treatment design.

Recent technological developments have enabled researchers to perform single-cell sequencing (SCS) experiments, where DNA from an individual cell is extracted, amplified, and sequenced. SCS provides high-resolution data for studying tumor evolution at unprecedented detail, e.g., it offers the possibility to identify branching topologies with high confidence or to solve the general problem of inferring the complete history of tumor evolution, even when all of the sequenced single-cells are extracted from a single-tumor biopsy sample. Early developments in inferring tumor evolutionary history from SCS data include probabilistic methods such as SCITE (Jahn et al. 2016), OncoNEM (Ross & Markowetz 2016), which employ the “infinite sites” assumption, and SiFit (Zafar et al. 2017), which relaxes this assumption by allowing violations at a given “cost”. More recently, SPhyR (El-Kebir 2018), a combinatorial optimization approach based on “Dollo” parsimony, and SiCloneFit (Zafar et al. 2019), which is an improved version of SiFit, were proposed. Finally, PhISCS-BnB (Sadeqi Azer et al. 2020), another combinatorially optimal tool utilizing a branch-and-bound strategy, and ScisTree (Wu 2020), a neighbor joining-based heuristic, are among the most recently developed methods that offer significant improvements in running time.

When both single-cell and bulk sequencing data of a tumor sample are available, two of the latest methods, namely B-SCITE (Malikic et al. 2019a) and PhISCS (Malikic et al. 2019b), can provide more accurate tumor phylogenies.

B-SCITE is a probabilistic method that integrates SCITE (Jahn et al. 2016) and CITUP (Malikic et al. 2015), whereas PhISCS is based on the use of combinatorial optimization and has two distinct implementations: PhISCS-I uses integer-linear programming while PhISCS-B formulates the same problem as a boolean constraint satisfaction program and solves it via available solvers for weighted max-SAT. Both approaches can also be applied to datasets consisting solely of SCS data.

As summarized above, available methods for tumor phylogeny reconstruction by the use of SCS data have important limitations. First, many of these methods employ Infinite Sites Assumption, ISA (even though some offer a provision for limited loss and concordant gain of mutations) and assume a uniform noise level (false negative as well as a false positive rate) - both subject to change with advances in our understanding of tumor evolution and SCS technology. More importantly, these methods aim to infer the most likely tumor phylogeny, and for that eliminate noise (due to, e.g., allele dropout or low sequence coverage) with a maximum likelihood/parsimony approach. As such, they all aim to solve an NP-hard problem from scratch, and thus cannot scale up to handle large SCS datasets. Even when the goal is to infer basic topological features of the tumor phylogeny rather than reconstructing it entirely, these methods cannot easily handle SCS data involving a few hundred mutations and cells. As a result, fast techniques for inferring key features of a tumor phylogeny, e.g., those that can discriminate linear from branching topologies, especially for SCS datasets with high noise levels (as per the current practice) are in high demand. Similarly, it is desirable to quickly infer whether any noise_elimination is necessary for constructing a perfect phylogeny. Finally, each of the existing tools has required a great deal of human effort in algorithmic design and implementation, as each technological advance in data generation necessitated the development of completely novel methodologies. It is thus highly desirable to have a general computational approach that can adapt to technological change, simply through training it with new data, without the need for explicit objective or noise profile modeling.

It may be possible to address these limitations via a *data-driven*, machine learning approach that considers a general set of functions and choose one that best fits a training dataset - which can be simulated or obtained through real-world measurements. Such an approach can, not only reduce the inaccuracies in noise profile modeling, but also identify implicit underlying patterns in the data or problem towards developing more realistic objectives. Recent advances in deep learning have demonstrated remarkable generalization of formulations for solving many problems (e.g., mastery over games such as Chess, Go (Silver et al. 2017), and Poker, or handling natural language tasks via BERT (Devlin et al. 2019) and most recently RoBERTa (Liu et al. 2019)), and it is possible that a single deep learning architecture may succeed in inferring distinct properties of tumor phylogenies when properly trained on a sufficient number of datasets.

In recent years, many computing applications have experienced a shift to data-driven approaches, from deciphering handwritten text (e.g., (Ciregan et al. 2012) for digit recognition) to natural language processing (NLP), e.g., (Devlin et al. 2019, Liu et al. 2019). Problems with poorly understood/formulated objectives such as those in structural biology (e.g., *AlphaFold* (Senior et al. 2020) for inferring the 3D structure of a protein sequence) seem to benefit considerably from deep learning methods. However, we are not aware of any data-driven approach for tumor phylogeny inference.

In this work, we offer the first data-driven tumor phylogeny reconstruction methods to address the limitations of existing strategies. We have used SCS data with deep neural networks and reinforcement learning to infer topological features of a tumor phylogeny, as well as the most likely evolutionary history of a tumor. In order to achieve this, we had to overcome several distinct challenges: (1) The neural network should ideally be designed so as to handle a varying number of cells and mutations. Alternatively, for models with fixed-sized inputs, it is desirable to use our domain knowledge to prepare the data in a manner that will facilitate success in predictions. (2) Given the use of neural networks, one needs a large number of samples for proper training. Unfortunately, the number of publicly available tumor SCS datasets are not sufficiently large to train deep learning models, thus we needed to produce a large number of simulated SCS datasets. (3) Errors/noise in SCS data add further complexity to the problem, and the proposed deep learning framework had to be evaluated in terms of noise tolerance. (4) The chosen architecture also expects a specific type of supervision which we must be able to compute. In order to perform noise reduction/elimination in the input *genotype matrix* (see below for a formal definition) extracted from SCS data, it is possible to offer supervision in the form of a dataset of noisy inputs along with their denoised versions. An alternative and cheaper supervision is offered by a feedback mechanism for determining whether a candidate output of the neural network is successfully denoised. A third alternative is given by a cost function that indirectly helps supervise a reinforcement learning process.

Inspired by novel deep learning approaches to combinatorial problems such as the *reinforce policy gradient algorithm* (Williams 1992) for the traveling salesperson (TSP) and the knapSack problems (Bello et al. 2017), and, to a limited degree the *NeuroSAT* approach (Selsam et al. 2019) for the satisfiability (SAT) problem using a single bit supervision, we established a computational framework to address all the above challenges as follows successfully. (i) We employed a reinforcement learning approach to train a denoising model without a need for the ground truth. The cost function we used is novel and problem-specific. (See Section S1.3). (ii) We devised a novel method to transform the input matrix (obtained from noisy SCS data) to feed to the neural network. This method incorporates the noise rates as well as coordinates and values of entries in the given matrix (See section S1.3.1). This representation also supports training and testing instances of different sizes and encourages robustness to permutation of rows and columns. (iii) We also present an example for incorporating domain knowledge to improve the performance of a machine learning model: this was achieved by sorting the columns of the input genotype matrix in a preprocessing step. (See Section S1.2.) (iv) The simulated input data we use with a given number of mutations and cells was obtained through a pipeline we developed - which along with the simulation tools from previous work, allows us to construct *unbiased* datasets to train our models.

### Results

We study three major problems.

First, given a noisy SCS dataset in the form of a genotype matrix, we consider the problem of inferring whether the most likely tumor phylogeny has linear topology or contains at least one branching event. The method we devised for this problem succeeds to learn and differentiate the two topology types in ~ 98% of randomly chosen noisy genotype matrices of size 100 × 100. The trained model runs at least 1000 times faster than the fastest available baseline techniques (see Table 2), which needs to first denoise the input and then examine the tree. The trained model is noise-tolerant, scalable, and can analyze real datasets (see Section 3.2). Moreover, having an efficient prediction function allows us to systematically study the relationship between the amount and type of noise in the input and the information preserved in the data to enable correct prediction.

Second, we present results on the problem of inferring whether a given binary matrix is conflict-free and thus admits a perfect phylogeny, or does not satisfy the so-called three-gametes rule. Our deep learning approach for this binary classification problem offers notably more accurate results in comparison to support vector machines. Crucially, sorting the columns of the input genotype matrix according to their binary representation as a preprocessing step further improves the accuracy of the trained models by 13-27% (See Table 3).

Third, we study the general problem of reconstructing the complete most likely tumor phylogeny. For solving this problem, we introduce a reinforcement learning based approach, which finds a solution for, e.g., 92% of the randomly generated noisy genotype matrices of size 10 × 10. Furthermore, our trained model on noisy genotype matrices of size 10 × 10 successfully solves a notable proportion of larger noisy genotype matrices, demonstrating the *generalizability* of our approach. See Table 1 for a summary of our approach’s key characteristics.

**Table 1:** A Summary of the key characteristics of our solutions for the problems defined in Section 2.

| Problems | The key characteristics of our Solutions |
| --- | --- |
| Branching Inference | • Based on a two-layer fully connected neural network<br>• Scalable<br>• Noise-tolerant<br>• Faster than available alternatives |
| Noise Inference | • Based on a two-layer fully connected neural network<br>• Has better accuracy compared to the support vector machines<br>• Its accuracy improves with our input preprocessing<br>• Provides insight about the capacity of machine learning models in studying key features of tumor evolution<br>• Does not provide improvement in running time over the available sophisticated combinatorial solution |
| Noise Elimination | • Based on a reinforcement learning framework<br>• Generalizes across matrix sizes, i.e., models trained on small matrices can solve larger matrices<br>• Typically slower than the existing methods |

## 2 Preliminaries

Given a positive integer *z*, let [*z*] denote the set {1, 2, *…*, *z*}. For a given matrix *M*, we let |*M*|_0_ denote the number of nonzero entries in *M*. For any boolean proposition P, we use Iverson bracket notation as

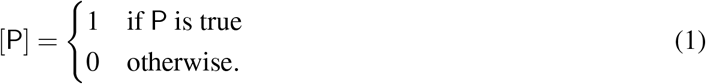

Assume that we have performed an SCS experiment where *n* single-cells, denoted by *C*_1_,*C*_2_, …,*C*_*n*_, were sequenced and *m* putative mutations, denoted by *M*_1_, *M*_2_, …, *M*_*m*_, were reported (in this work, we focus on single-nucleotide variants at loci that are homozygous in normal cells). We define a *genotype* of cell *C _i_* as a binary vector of length *m* having *j*-th coordinate equal to 1 if and only if *C*_*i*_ harbors mutation *M*_*j*_. We distinguish between the *true* and *noisy* genotypes of a cell, where the latter is obtained from SCS data and typically contains some false mutation calls.

We can combine true genotypes of the sequenced cells into a binary matrix *A* having *n* rows (corresponding to the cells) and *m* columns (corresponding to the mutations). More formally, *A*[*i*, *j*] = 1 if and only if cell *C*_*i*_ harbors mutation *M*_*j*_. Similarly, we define matrix *A*′ by combining noisy genotypes. In other words, *A*′[*i*, *j*] = 1 if and only if, after single-cell data processing and mutation calling steps, mutation *M*_*j*_ was reported to be present in cell *C*_*i*_. For the convenience of notation, below, we will mostly use *a*_*i*, *j*_ and *a*′_*i,j*_ assuming that they equal to *A*[*i*, *j*] and *A*′[*i, j*], respectively.

Single-cell genotype matrix *D* (of dimension *n* × *m* and with terms denoted by *d*_*i, j*_) is said to represent a *perfect phylogeny* if it satisfies the *three-gametes* rule, i.e.,

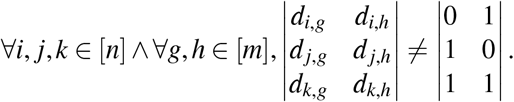

Notice that the rows and columns in the definition above can be in any order. In the case of equality, the row-triplet (*i*, *j*, *k*) and column-pair (*g*, *h*) is said to have a *conflict* or a *three-gametes rule violation*. We also refer to a matrix without such conflict as a *conflict-free matrix*. It is well known that, under the infinite sites assumption, for any conflict-free matrix, there exists a phylogenetic tree depicting the evolutionary history of the tumor such that: (i) the set of mutations assigned to the edges of the tree equals the set of mutations given by the matrix (ii) each mutation appears exactly once in the tree (iii) each of the single-cells represents exactly one leaf of the tree (iv) set of mutations occurring on the path from the root to a given single-cell (i.e., leaf) equals to the set of mutations present in the cell (this set is given by the related row of the matrix) (Gusfield 1997, Malikic et a l. 2019b, Edrisi et al. 2019). In addition, it is trivial to show that, under the infinite sites assumption, any noise-free single-cell genotype matrix is conflict-free. Therefore, instead of searching for the most likely tree of tumor evolution directly in the space of trees (as was done, for example, in (Jahn et al. 2016) and (Malikic et al. 2019a)), we can first search in the space of conflict-free matrices (as was done, for example, in (Malikic et al. 2019b)) and then, using the algorithm described in (Gusfield 1997), convert the obtained matrix into a tree. Note that in this work, we also consider *clonal trees* of tumor evolution, typically obtained from phylogenetic trees by discarding the attachment of single-cells as leaves. We refer to the section “Tree models of tumour evolution” in (Malikic et al. 2019a) for a detailed definition of the clonal tree.

To generate conflict-free matrices representing instances of *A*, we used the most recent version of the ms simulator, which was first introduced in (Hudson 2002) and has been used in earlier studies on phylogeny inference from bulk or single-cell sequencing data (Bonizzoni et al. 2018, El-Kebir 2018). To obtain instances of *A*′ which are not conflict-free, we added noise to *A* by assuming an independent and identically distributed (i.i.d.) noise process across the mutated loci (similarly as was done in most of the available studies, e.g., (Jahn et al. 2016, Ross & Markowetz 2016, Malikic et al. 2019a,b)). More precisely, we choose to *flip* each entry independently with a probability depending on the value of the entry and predefined false positive (*α*) and false negative (*β*) rates. To ensure the correctness of the labels and get a fixed number of instances that are not conflict-free, we check each generated matrix *A*′ to verify that it contains at least one conflict. If it does not, we repeat the above procedure until a matrix *A*′ containing one or more conflicts is obtained. See Supplementary Section S3 for details related to the use of ms and generation of matrices *A*′.

Under the assumption of independent and identically distributed (i.i.d.) noise process as mentioned above, the conditional probability that a conflict-free matrix *B* (with entries denoted by *b*_*i*_, _*j*_) generates *A*′ is:

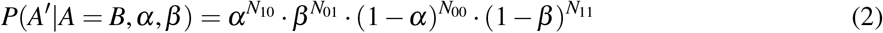

where *N*_*pq*_ for each pair (*p*, *q*) ∈ {(0, 0),(0, 1),(1, 0),(1, 1)} is defined as

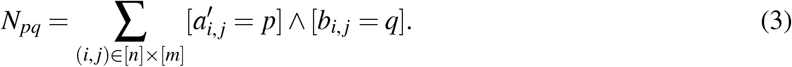

The detailed derivation of the above formula can be found in Equations (2)-(3) in (Malikic et al. 2019b). Note that *P*(*A*′|*A* = *B*, *α*, *β*) represents the likelihood that *B* is the true single-cell genotype matrix, assuming that noisy genotype matrix *A*′ was obtained from SCS data (or, shortly, the *likelihood of B*, as will be used throughout the rest of the manuscript).

In the remainder of this section, we will provide a formal definition and brief overview of each of the three problems considered in this work. Then, in Sections S1.1, S1.2, and S1.3, solutions to each of these problems are discussed in more detail.

### 2.1 Branching Inference Problem

As mentioned in Section 1, the problem of inferring whether a tumor contains divergent subclones evolving through distinct branches of its phylogeny has potential applications in improving cancer treatment. This motivated us to consider the following computational problem.

#### no_branching problem

Given a noisy binary matrix *A*′ obtained as the output (after mutation calling step) of a single-cell sequencing experiment, decide whether the most likely clonal tree of tumor evolution of the sequenced tumor has a linear topology or a topology which contains at least one branching event.

To solve the above problem, instead of inferring the complete most likely clonal tree of tumor evolution and then checking whether it has linear topology or not, here we obtain equivalent results by working solely in the space of binary matrices. This is possible since the set of conflict-free matrices, which imply linear topology, can be precisely characterized. Before providing the formal mathematical characterization of this set, we introduce the set of *staircase matrices*. A staircase matrix is any binary matrix *S* such that (i) *S*[*i*, *j*] ≤ *S*[*i* + 1, *j*] for all pairs of integers (*i*, *j*) such that 1 ≤ *i* ≤ *n* − 1 and 1 ≤ *j* ≤ *m*, where *n* and *m* respectively denote the number of rows and columns of *S*, and (ii) *S*[*i*, *j*] ≤ *S*[*i*, *j* + 1] for all pairs of integers (*i*, *j*) such that 1 ≤ *i* ≤ *n* and 1 ≤ *j* ≤ *m* − 1. It can be proved that a binary matrix *D* is conflict-free and implies a clonal tree with linear topology if and only if its columns and its rows can be reordered so that the newly obtained matrix belongs to the set of staircase matrices.^1^ In other words, the set of conflict-free matrices that imply trees having linear topology consists exactly of all binary matrices that can be converted to staircase matrices by reordering their rows and columns. In solving no_branching problem, we extensively rely on this fact.

We say that any conflict-free binary matrix *D* which implies a clonal tree with linear topology has the *nobranching* property. Given the above, no_branching problem can also be defined as the problem of inferring whether the most likely denoised version of a given binary matrix *A*′ has no-branching property.

It was very recently shown that no_branching problem is NP-hard even when a false positive rate *α* of single-cell data is set to 0.^2^

We study this problem in the machine learning framework introduced as follows: Given a conflict-free matrix *D*, we define a function Sing(*D*) : {0, 1}^*n*×*m*^ → {0, 1} which takes value 1 if *D* has the no-branching property, i.e., implies a linear topology, and 0 otherwise. Our goal for the no_branching problem is to choose a function *f* (*D*) : {0, 1}^*n*×*m*^ → {0, 1} from a domain of functions described in Section S1.1, such that *f* approximates Sing with the highest possible accuracy, where the accuracy of *f* is defined as the proportion of potential input matrices *D* where *f* (*D*) = Sing(*D*).

### 2.2 Noise Inference Problem

In the noise_inference problem, our goal is to investigate whether neural network models can help us to successfully check whether a given binary matrix admits perfect phylogeny, which is equivalent to checking whether the matrix contains at least one conflict. So we consider the following problem:

#### noise_inference problem

Given a binary matrix *A*′, decide whether *A*′ is conflict-free.

This problem is well-studied in literature and has a linear time algorithm (Gusfield 1991). Here, we study the capacity of machine learning models to train to solve this problem. We start by imposing the machine learning framework on the noise_inference problem as follows: Let the function Icf(*A*′) : {0, 1} ^*n*×*m*^ → {0, 1} have value 1 if *A*′ contains at least one conflict, and 0 if it is conflict-free. Our goal for the noise_inference problem is to choose a function *f* (*A*′) : {0, 1}*n*×*m* → {0, 1} from a domain of functions described in Section S1.2, such that *f* approximates Icf with the highest possible accuracy, where this time, the accuracy of *f* is defined as the proportion of potential input matrices *A*′ where *f* (*A*′) = Icf(*A*′).

### 2.3 Noise Elimination Problem

The problem of denoising a binary matrix obtained from SCS data to obtain a conflict-free matrix has recently attracted attention in the field of tumor phylogenetics (Ciccolella et al. 2018, Edrisi et al. 2019, El-Kebir 2018, Malikic et al. 2019b, Sadeqi Azer et al. 2020). The general denoising problem, which we call the noise_elimination problem, is defined as follows:

#### noise_elimination problem

Given the input consisting of noisy genotype matrix *A*′ and noise parameters *α* and *β*, our objective is to find conflict-free matrix *B* such that the likelihood of *B*, i.e., the probability *P*(*A*′ | *B*, *α*, *β*), is (ideally) maximized.

The noise_elimination problem is the most computationally challenging problem considered in this paper since it cannot be easily formulated as a classification, regression, or clustering problem, which are better suited for deep learning approaches. More importantly, the problem is NP-hard (Chen et al. 2006) and available principled solvers deploy sophisticated methods such as mixed-integer linear programming, constraint satisfaction programming, Markov chain Monte Carlo towards its solution for practical instances (Chen et al. 2006, Jahn et al. 2016, Malikic et al. 2019a,b). Finally, since the problem is computationally intractable, it is not possible to construct large matrices with known optimal solutions to be used for training a neural network. This issue alone makes it impossible to use several successful machine learning techniques for our purposes.

## 3 Experiments

In this section, we discuss our experimental setup and results. Additional details related to hardware specifications, the generation of simulated data, and hyperparameter tuning are presented in Supplementary Sections S2, S3, and S4 to ensure reproducibility.

### 3.1 Experimental Setup

All experiments presented here were performed using deep learning (DL) nodes in *Carbonate*, a computation cluster at Indiana University (Stewart et al. 2017). In order to ensure the reproducibility of our results, we provide the specifications of these nodes, along with the version numbers of the packages used, in the README.md file of our repository.

We note that the accuracy values in Figures 1, 3, and 4 are higher for the evaluation phase than the training phase. This reflects the fact that the error during training was measured in a noisy manner using dropout, which is necessarily worse than the true error rate of the network.

**Figure 1:**
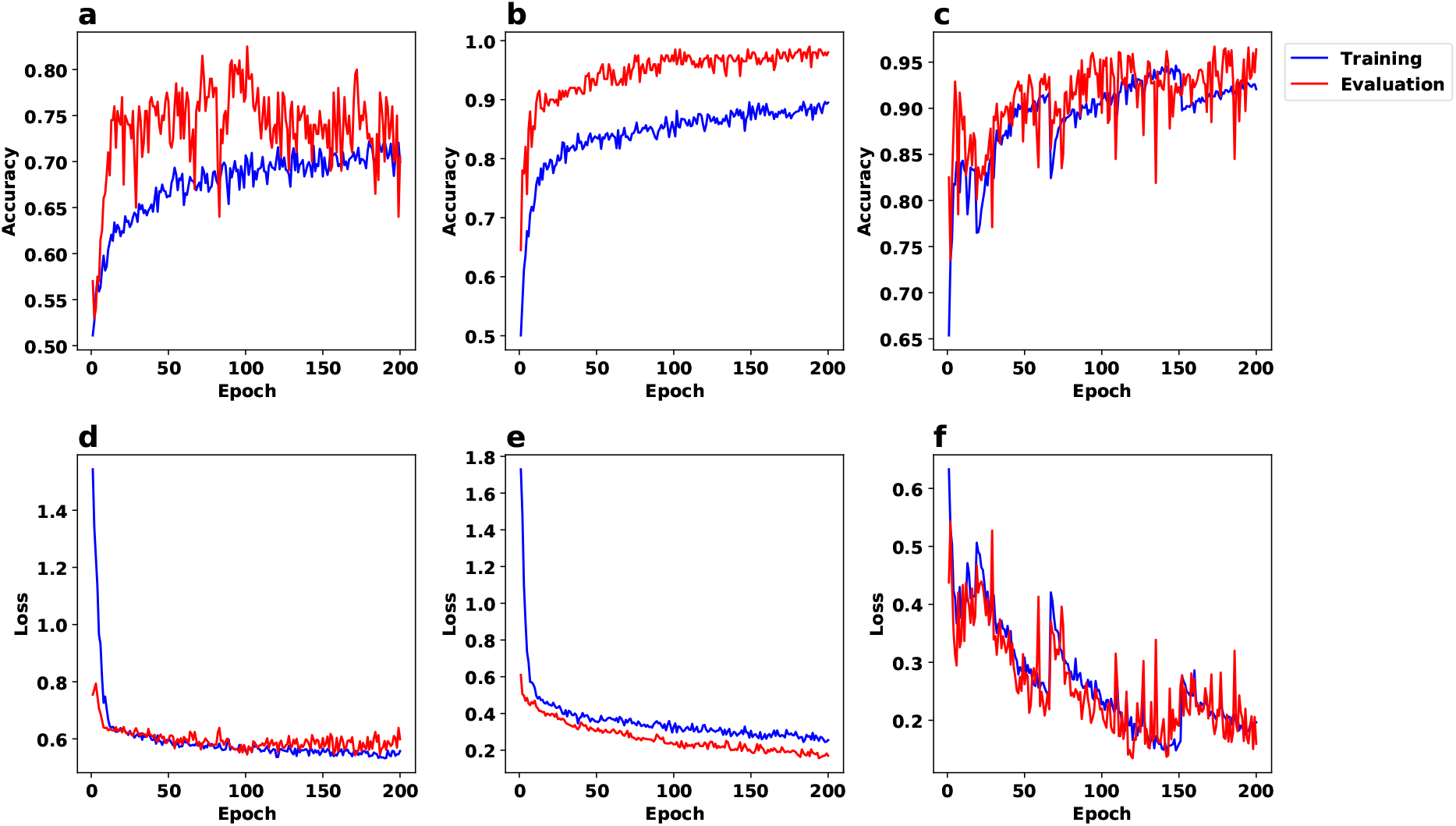
The accuracy (first row) and binary cross entropy loss (Bishop 2006) (second row) plots corresponding to the results for no_branching problem and the architecture described in Section S1.1. We used a dataset of 2000 matrices with dimension 100 × 100 and hidden layer of size 10 in (a,d), the same dataset as in (a,d) with hidden layer of size 100 in (b,e), and a dataset of 10000 matrices with dimension 500 × 500 and hidden layer of size 100 in (c,f).

### 3.2 Branching Inference

Recall the no_branching problem, where the goal is to identify whether the tumor phylogeny has a linear or a branching topology.

#### Learning curves

In the first set of experiments for this problem, we split the dataset to training and evaluation sets with a ratio of 9 to 1. We designed several experiments with different settings and all of them show success in predictions. We include results for three settings in Figure 1: (a,d) a dataset of 2000 matrices of size 100 × 100 and a hidden layer of size 10, (b,e) same dataset, now with a hidden layer of size 100, and (c,f) a dataset of 10000 matrices of size 500 × 500 and hidden layer of size 100. In each of these experiments, we run the training process for 200 epochs. Note that the model distinguishes up to 98% of randomly chosen instances of size 100 × 100 correctly - see panel (b) in Figure 1.

#### Comparison of Running Time for the no_branching problem

We performed an experiment to compare the running time of our approach and recently published method PhISCS (Malikic et al. 2019b), which we use as a baseline for the no_branching problem.

In order to assess the performance improvement obtained by our approach, we ran it on two simulated genotype matrices, one 100 × 100 and another 500 × 500. Each matrix was then subjected to random noise (*α* = 0.002 and *β* = 0.2) 50 times before we applied our approach; we reported the worst running time of our approach out of these 50 runs.

PhISCS first denoises the input matrix before it builds the tree, based on which we can check whether its topology is linear or not. Based on our experience with running PhISCS, its running time depends heavily on the noise levels, i.e., the higher the noise level, the longer its running time. Therefore, we decided to run PhISCS on the noise-free versions of these two input genotype matrices. Despite this disadvantageous setting for our approach, it turned out to be more than 1000 times faster than PhISCS, as can be seen in Table 2. Note that, since the running time of PhISCS was prohibitive for the larger matrices, we could not conduct a more comprehensive study of its output characteristics, including its accuracy.

**Table 2:** A comparison of the running times (measured in seconds) between our approach and the baseline for the no_branching problem. For our approach, we tested 50 noisy cases (*α* = 0.002, *β* = 0.2) for both matrix sizes, and report the maximum running time we observed. For the baseline, we report the best-case running time for noise-free input matrices (adding noise increases the running time substantially).

| Input Matrix Size | $100 \times 100$ | $500 \times 500$ |
| --- | --- | --- |
| Time: Our approach | 0.0372 | 0.0537 |
| Time: PhISCS | $> 80$ | $> 3772$ |

#### Evaluation of noise-tolerance for the no_branching problem

In another experiment, we tested the noise tolerance of our model on five different values for *β/α*, i.e., the false negative to false positive rate ratio. For each value in Figure 2, we plotted the percentage of inputs whose topologies are correctly distinguished as a function of the false negative rate *β*. We observe that when the false positive value is relatively low, e.g., in case where *α* = 0.03, *β* = 0.3 in Figure 2 panels (c,d,e), the accuracy of the model remains close to perfect across all (realistic) values of the false negative rate *β*. Notice that as *β/α* increases in panels (c,d,e), the accuracy increases.

**Figure 2:**
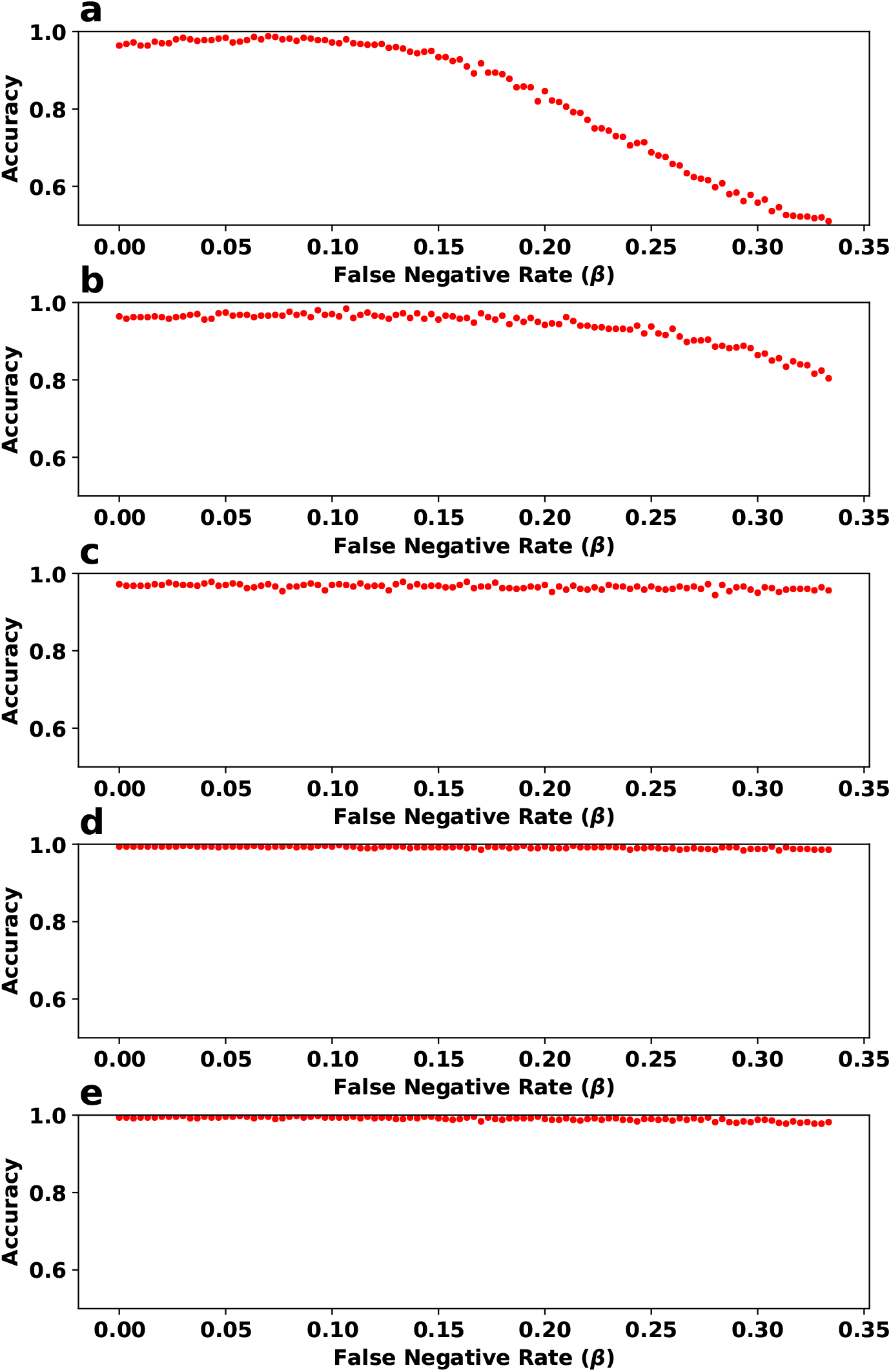
The evaluation of noise tolerance for our model for the no_branching problem. In each panel, the y-axis (accuracy) represents the fraction of instances classified correctly from 500 randomly chosen instances. These instances are not seen during training and consist of an equal number of instances from each topology type (See Section S1.1). The trained model consists of two layers with size 100 and 0.9 drop out rate. This model was trained using 2000 matrices of size 100 × 100 with <u>no noise</u> for 200 epochs. The five panels differ in the ratio of false negative to false positive rates: (a) *α* = *β*, (b) *α* = *β/*2, (c) *α* = *β/*10, (d) *α* = *β/*100 and (e) *α* = *β/*1000. Note that the noise tolerance of the method gets more robust as the relative value of *α* decreases.

Note that the efficiency of our approach as shown in Table 2 enabled us to run an extensive number of experiments to obtain the data points in Figure 2. The plots in this figure suggest a new understanding of the relationship between the amount of preserved information that can be captured by our neural network in the presence of different noise levels. For example, for settings where false negative and false positive rates are equal, a false negative rate of ≤ 0.15 preserves the deduced topological outlook fairly consistently. The drop in accuracy after this point seems steeper.

### 3.3 Noise Inference

We trained and evaluated our proposed neural network for the noise_inference problem introduced in Section S1.2, on multiple datasets. We consider two sizes for the input matrices: 10 × 10 and 25 × 25. For each size, we experimented with two distinct datasets that differed with respect to the amount of noise introduced. Note that in the context of this problem, unlike both the no_branching problem and the noise_elimination problem, the lower the noise level, the harder the decision problem becomes. In order to assess the full capabilities of our model, we used lower values for *α* and *β* than typical false positive and false negative rates observed in real SCS datasets. (As mentioned in Section 2, even at lower noise levels, we have ensured that the noisy matrices include at least one conflict.)

In Figure 3, panels (a,c) show accuracy and binary cross entropy loss (Bishop 2006) for training and evaluation datasets, respectively comprised of 2*M* and 2*K* matrices, each of size 10 × 10, where half of each set is noisy with *α* = 0.002 and *β* = 0.1 (with the remaining half being conflict-free). Panels b and d show results for *α* = 4 × 10^−4^ and *β* = 0.02 (again accompanied by an equal number of conflict-free matrices, both in training and evaluation). We observe an accuracy of 90% in the dataset with a higher noise level. This confirms that distinguishing matrices that contain at least a conflict from those without any conflict gets harder as the noise level decreases. It is worth mentioning that the noise levels observed in practice are higher than these values.

**Figure 3:**
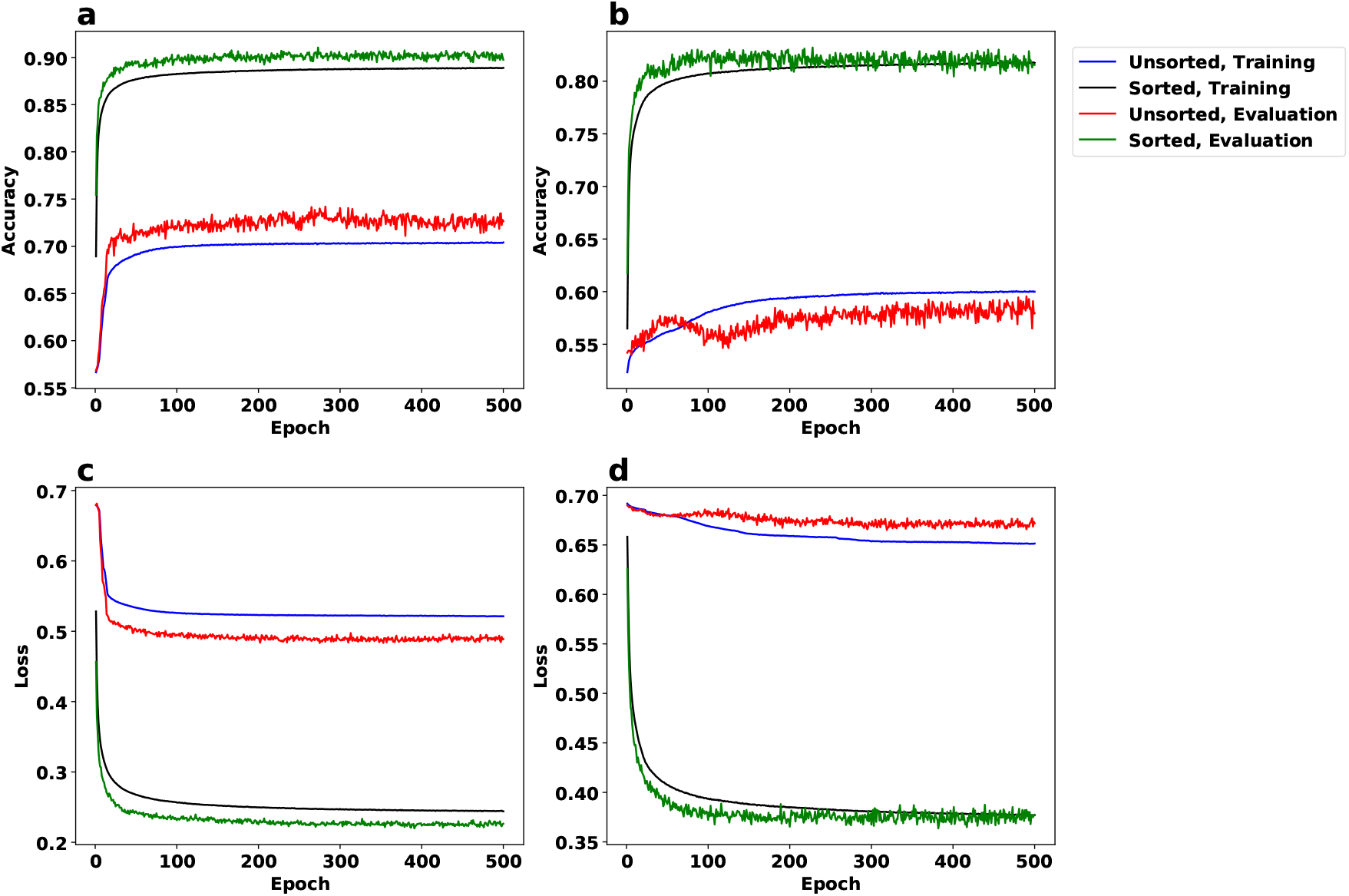
The accuracy and loss (Bishop 2006) plots for the noise_inference problem using the neural network described in Section S1.2. The datasets in these experiments include matrices of size <u>10 × 10</u>. The colors of the plots correspond to whether column-wise sorted or unsorted datasets were used - as well as whether the plot corresponds to the training phase or evaluation. We use *α* = 0.002 and *β* = 0.1 in panels (a,c) and *α* = 4 × 10^−4^ and *β* = 0.02 in panels (b,d). Note that panels (a,b) depict accuracy whereas panels (c,d) depict binary cross entropy loss (Bishop 2006).

Results for matrices of size 25 × 25 are shown in Figure 4.

**Figure 4:**
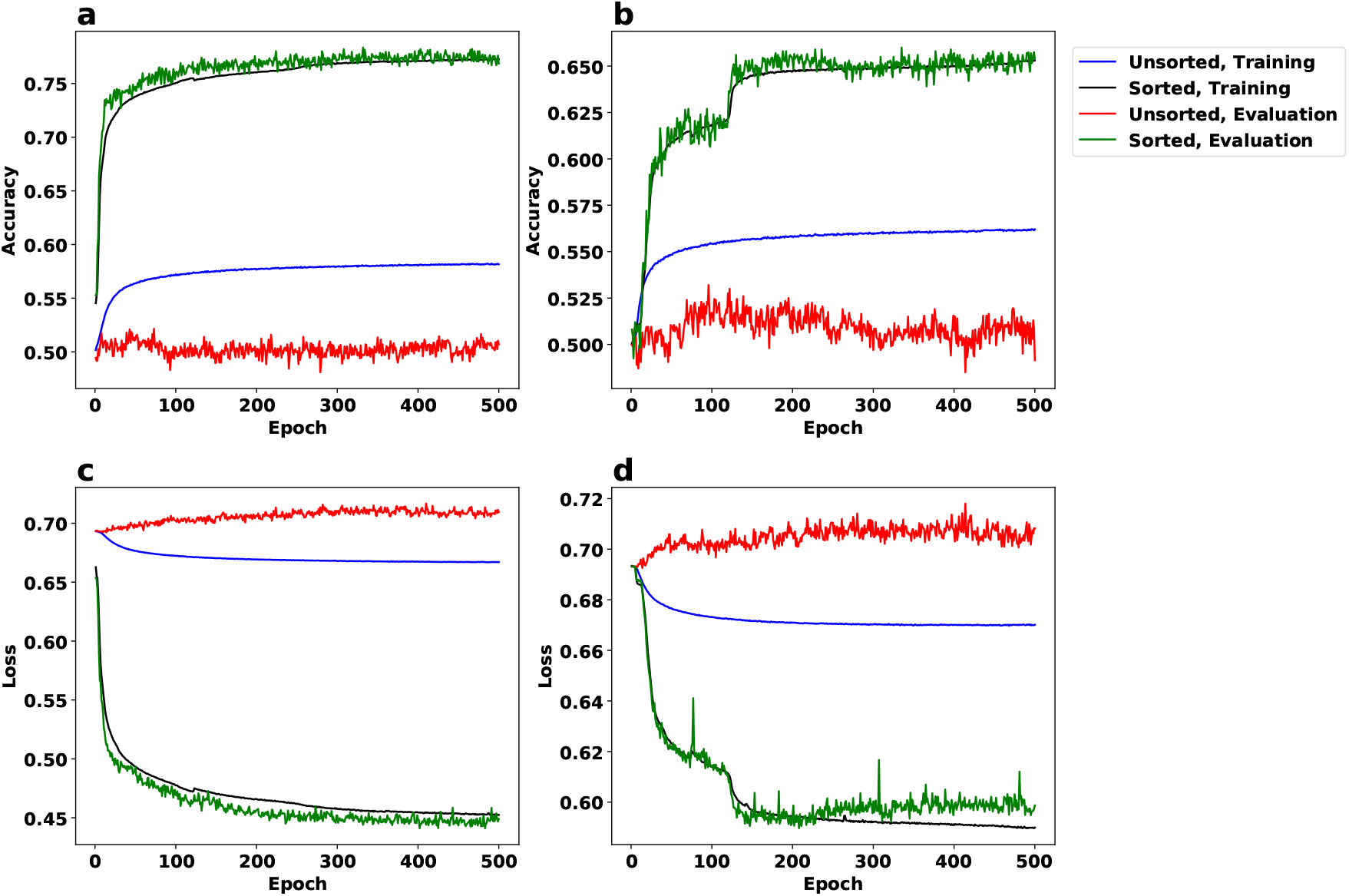
The accuracy and loss (Bishop 2006) plots for the noise_inference problem using the architecture described in Section S1.2 on 25 × 25 matrices. The plots for column-wise sorted and unsorted genotype matrices, as well as those for training and evaluation phases, are depicted in distinct colors. The noise settings for this figure are *α* = 3.2 × 10^−4^ and *β* = 0.016 for panels (a,c), and *α* = 6.4 × 10^−5^ and *β* = 0.0032 for panels (b,d). Note that panels (a,b) depict accuracy, while panels (c,d) depict binary cross entropy loss (Bishop 2006).

We used *α* = 3.2 × 10^−4^and *β* = 0.016 in panels (a,c) and *α* = 4 × 10^−4^ and *β* = 0.02 in panels (b,d). As mentioned in Section S1.2, considering columns of the input genotype matrix as binary vectors and sorting them in the non-decreasing order as a preprocessing step is expected to improve the prediction accuracy. As summarized in Table 3 this is well supported by the results presented in Figure 3 and Figure 4. As shown in Table 3, the accuracy of our approach improves if the input genotype matrix is sorted by its columns. This can be partially explained by the fact that sorting improves the locality of reference for mutations and makes it easier for the neural network to identify conflicts in a way reminiscent to the preprocessing step of the algorithm proposed by Gusfield (Gusfield 1991).

**Table 3:** The impact of the preprocessing the input genotype matrix by sorting its columns as described in Section S1.2 on the evaluation accuracy for the noise_inference problem. The accuracy values for unsorted (fourth column) and column-sorted genotype matrices (fifth column) are reported as percentages rounded to the nearest integer.

| Input Matrix Size | $\alpha$ | $\beta$ | Unsorted Acc. | Sorted Acc. | Figure |
| --- | --- | --- | --- | --- | --- |
| $10 \times 10$ | 0.002 | 0.1 | 72 | 90 | Fig. 3 (a) |
| $10 \times 10$ | $4 \times 10^{-4}$ | 0.02 | 60 | 81 | Fig. 3 (b) |
| $25 \times 25$ | $3.2 \times 10^{-4}$ | 0.016 | 50 | 77 | Fig. 4 (a) |
| $25 \times 25$ | $6.4 \times 10^{-5}$ | 0.0032 | 52 | 65 | Fig. 4 (b) |

In order to evaluate the relative advantage of neural networks against other machine learning techniques for the noise_inference problem, we explored the use of support vector machines (SVM) (Bishop 2006) for the same task. Figure 5 shows the training and evaluation accuracy for (column-sorted) 10 × 10 matrices using SVM.

**Figure 5:**
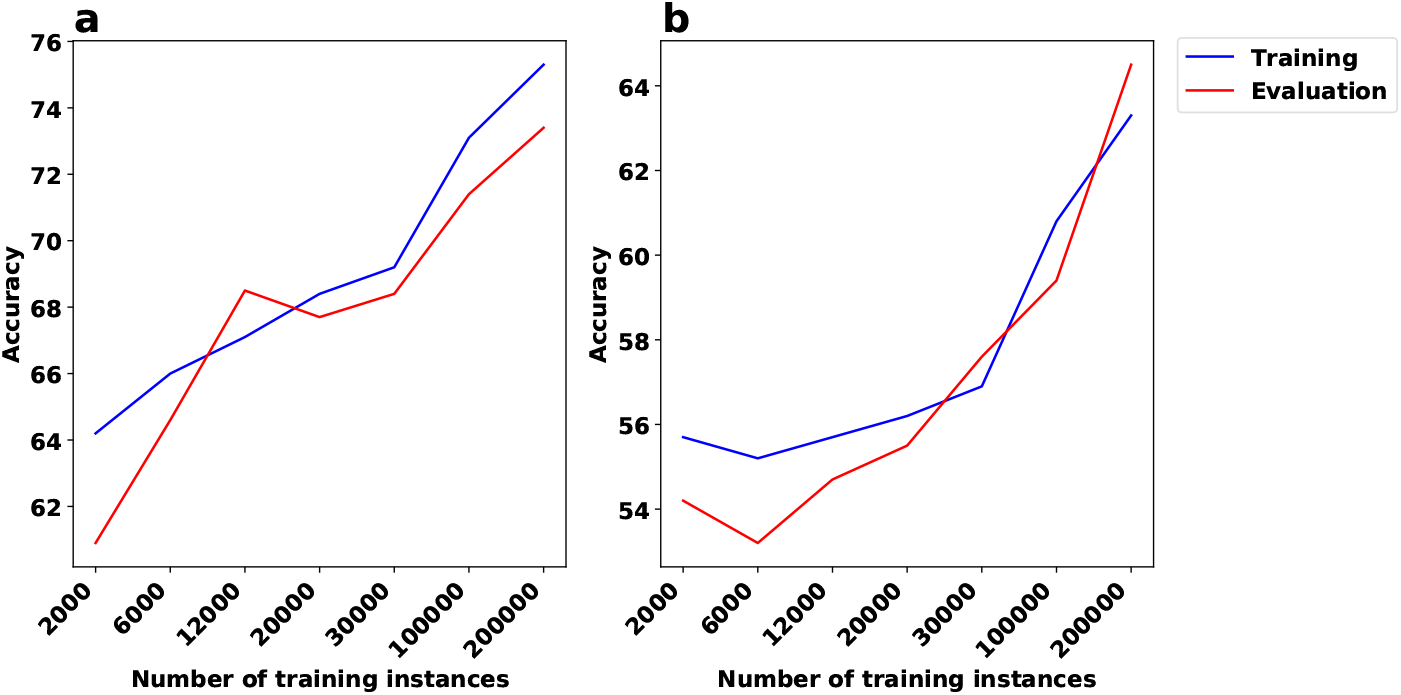
The training and evaluation accuracy in the noise_inference problem for SVM. All matrices for this experiment are of size 10 × 10 and are column-sorted. The x-axis corresponds to the training sample size. (a) Training and evaluation accuracy for *α* = 0.002 and *β* = 0.1 (for the noisy matrices). (b) Training and evaluation accuracy for *α* = 4 × 10^−4^ and *β* = 0.02 (for the noisy matrices).

The x-axis in this figure represents the training sample size. Since SVM is not memory efficient, the maximum number of input instances that can be used in training is limited by our computational resources. That is why the training procedure is stopped before the accuracy plot has converged. We use radial basis function (RBF) (Bishop 2006) as the kernel in all of our experiments with SVM. RBF is the most commonly used kernel in practice. In our experiments, it consistently showed better performance than other kernels (e.g., polynomial).

Observe that the accuracy of the neural network model on the evaluation data is notably higher than that for SVM. E.g. the final evaluation accuracy values in Figure 3 vs. Figure 5 is 90% vs 73% in the first setting and 81% vs 64% in the second.

### 3.4 Noise Elimination

We train the network shown in Figure S1 using the reinforcement learning framework described in Section S1.3 on matrices of size 10 × 10, with *α* = 4 × 10^−4^ and *β* = 0.02. We constructed an evaluation dataset of 100 matrices with the *α* and *β* values identical to that of the training dataset; although this noise profile is not currently observed in single-cell sequencing datasets, they have been achieved by emerging single clone sequencing datasets (Pérez-Guijarro et al. 2020).

We made sure that the training and evaluation datasets had no overlaps. Our trained model computed a conflict-free output matrix for 92% of the evaluation matrices.

Let *o*_*FP*_, and *o*_*FN*_ respectively denote the number of flips from 0 to 1 and from 1 to 0 that we introduced to a ground truth conflict-free matrix *A* to produce *A*′. Similarly, let *r*_*FP*_ and *r*_*FN*_ respectively denote the number of flips from 1 to 0 and 0 to 1 made by the model on the noisy input matrix *A*′ in order to produce a conflict-free output matrix *B*. Figure 6.a shows the histogram of *r*_*FP*_ − *o*_*FP*_ (left side) and *r*_*FN*_ − *o*_*FN*_ (right side) for the 92 instances of the problem for which our model produced a conflict-free matrix. Importantly, in the majority of cases, *r*_*FP*_ ≤ *o*_*FP*_ and *r*_*FN*_ ≤ *o*_*FN*_, indicating that the number of flips used by our approach is at most that of the number of flips introduced as noise (in some of the cases, there are solutions that result in a conflict-free with fewer flips than that added as noise). We expand this visualization further in Figure 6.b using a heat map.

**Figure 6:**
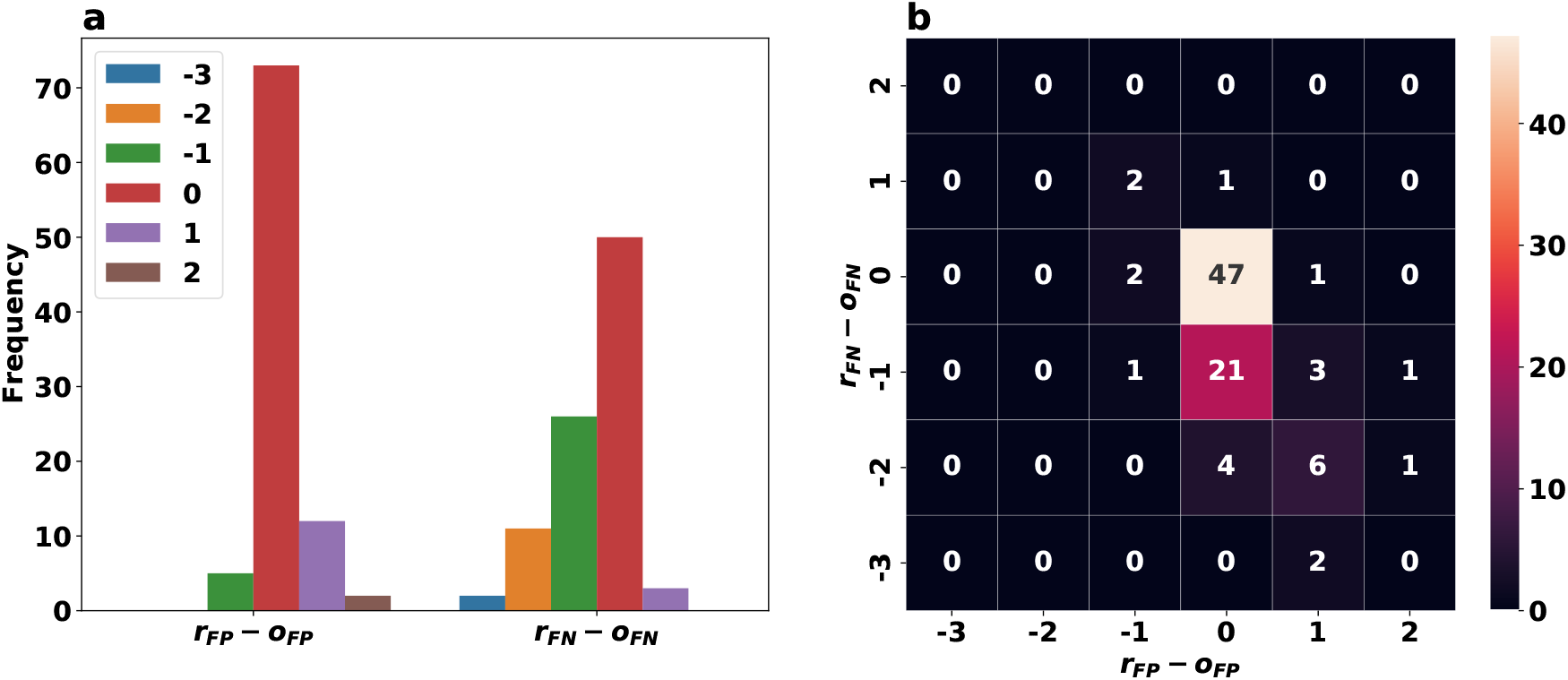
The difference between the number of flips made by our approach and the flips introduced as noise by the simulation, i.e., to the ground truth to obtain the input genotype matrices, given in histograms in panel (a) and a heat map in panel (b), for the noise_elimination problem. The left histogram in panel (a) and the x-axis in panel (b) correspond to false positives. Similarly, the right histogram in panel (a) and the y-axis in panel (b) correspond to false negatives. Lower values for *r*_*FN*_ − *o*_*FN*_ and *r*_*FP*_ − *o*_*FP*_ imply a good performance for our approach. The matrices used in this experiment are of size 10 × 10 and the noise rates are *α* = 4 × 10^−4^ and *β* = 0.02.

In order to quantify how well our approach′s distribution of flip types compare to those flips introduced as noise, we introduce ratio_rl_ = *r_FN_/*(*r*_*FP*_ + *r*_*FN*_) and ratio_original_ = *o_FN_/*(*o*_*FP*_ + *o*_*FN*_). The x and y-axis in Figure 7 represents ratio_rl_ and ratio_original_, respectively. Observe that most of the evaluation cases are on the line ratio_rl_ = ratio_original_.

**Figure 7:**
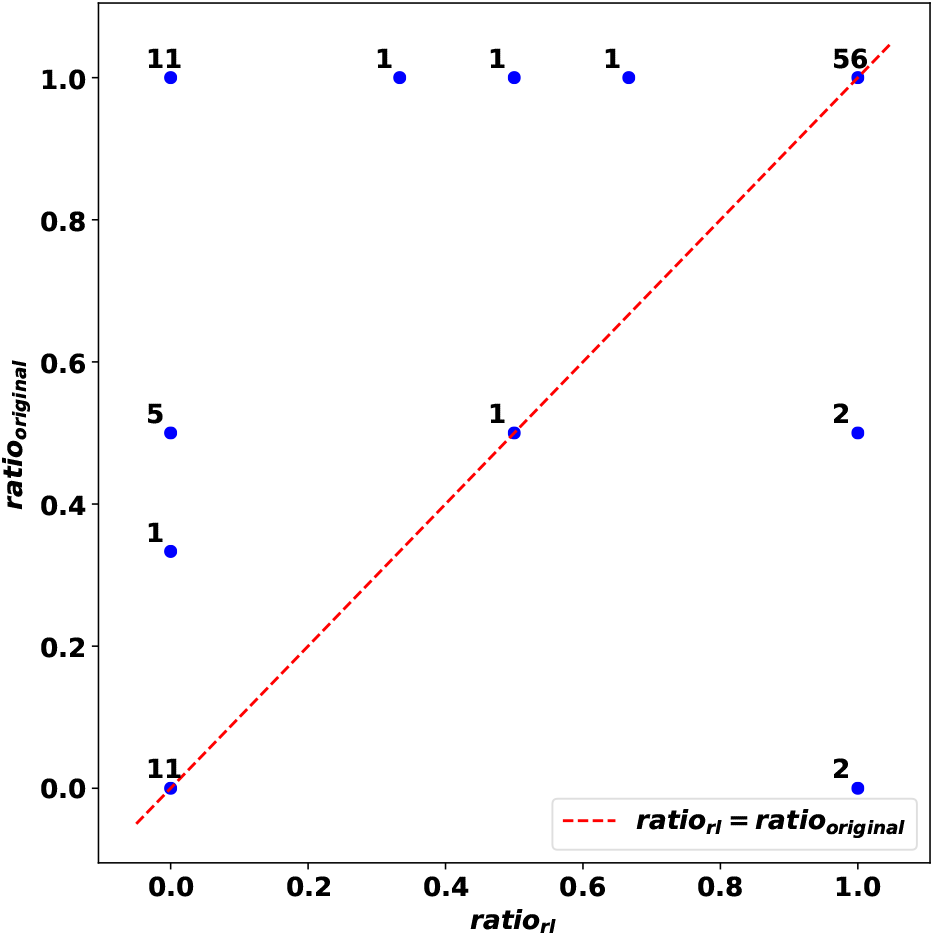
A quantified visualization of how flips made by our approach correspond to those added as noise for the noise_elimination problem. The plot compares the ratio_rl_ with ratio_original_ for matrices of size 10 × 10. Each blue dot is annotated by the number of respective evaluation cases. The red dotted line represents ratio_rl_ = ratio_original_. The closeness of the blue dots for the majority of the evaluation cases to the red dotted line implies the recovery of the ratio of introduced flip types by our approach.

Next, we compare trees implied by the conflict-free matrices reported by our reinforcement learning approach with the trees obtained by running PhISCS (Malikic et al. 2019b). As input, we use the same dataset that was used for experiments related to Figures 6 and 7. In all comparisons, we use recently developed Multi-labeled Tree Similarity Measure (MLTSM) for comparing trees of tumor evolution (Karpov et al. 2019). As shown in Figure 8, our approach achieves a tree reconstruction accuracy similar to PhISCS. It is worth noting that, on average, the reinforcement learning approach and PhISCS take 36.804 and 0.142 seconds, respectively, to process each matrix.

**Figure 8:**
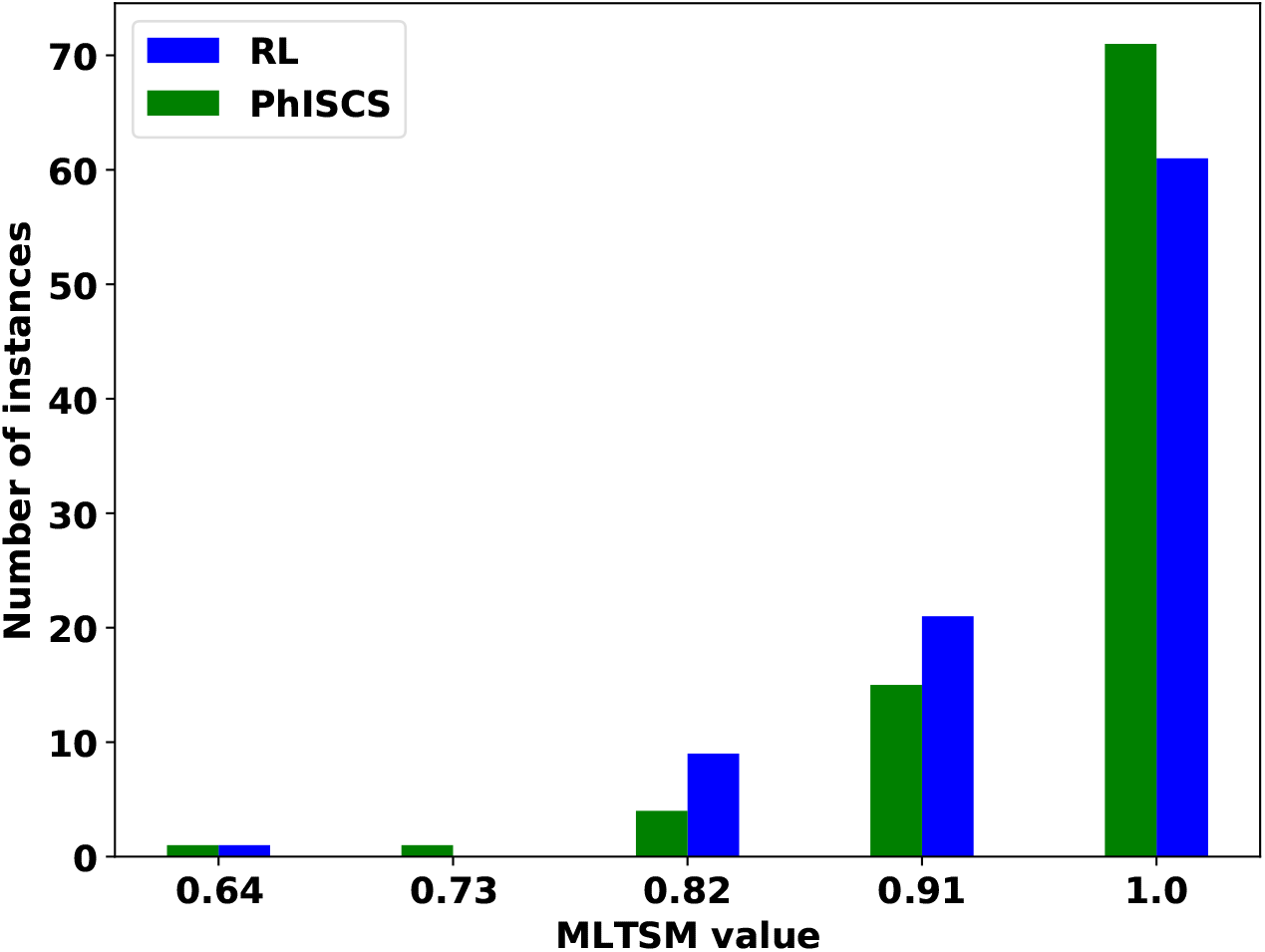
A comparison of the trees reconstructed using our reinforcement learning approach and the trees reported by PhISCS. MLTSM tree similarity measure is used in all comparisons. The y-axis shows the number of instances for which MLTSM value shown on the x-axis is obtained. The comparison is performed on the set of 92 matrices of size 10 × 10 with the false positive and false negative rates equal to 4 × 10^−4^ and 0.02, respectively.

Finally, we evaluated how well our trained models generalize to input matrices with varying dimensionality. In Figure 9 we used a model trained on 10 × 10 matrices for this purpose. Observe that this trained model can find a conflict-free matrix for a notable fraction of matrices with larger dimensionality. Note that the model found these conflict-free matrices by flipping at most 3 coordinates in each noisy matrix. As expected, our success rate decreases when the dimensionality increases.

**Figure 9:**
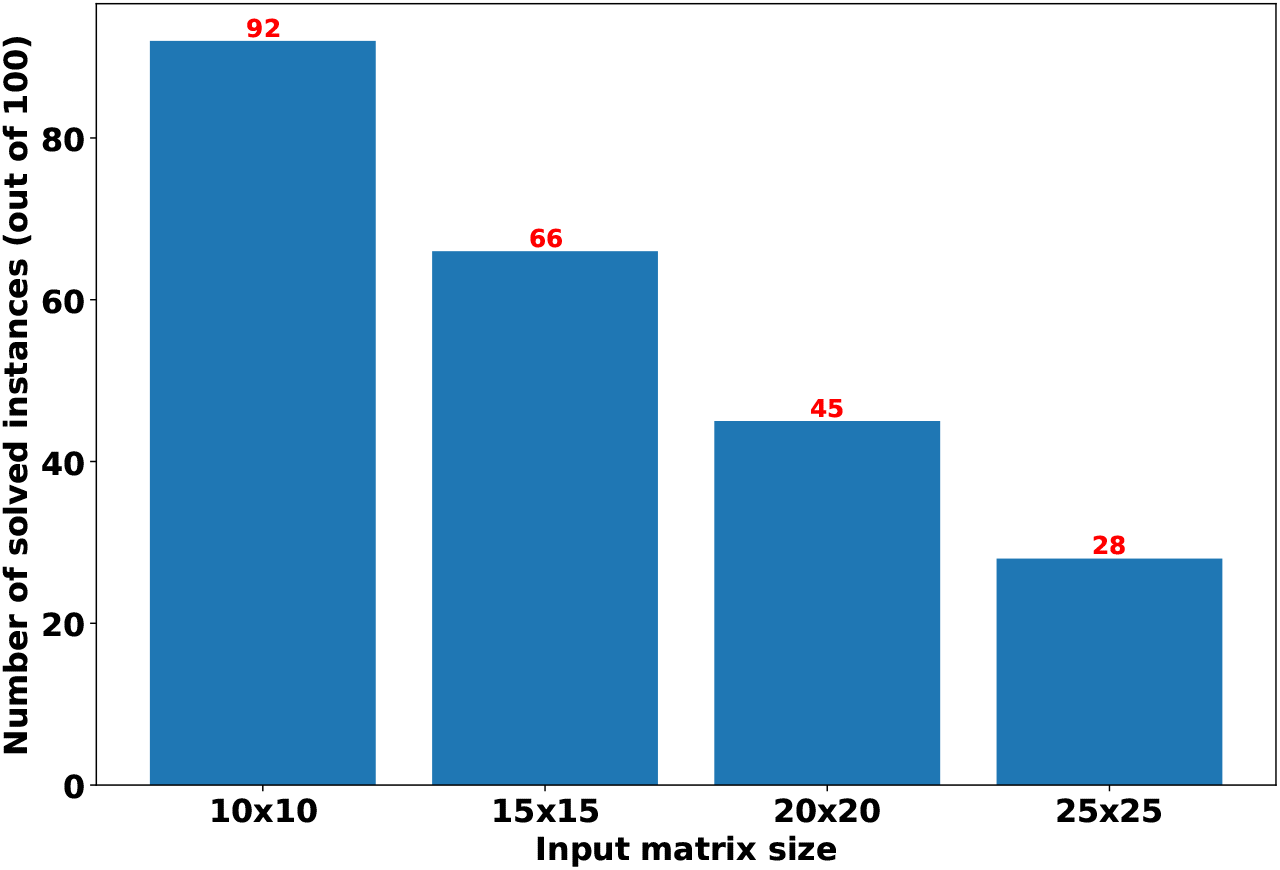
The evaluation of generalizability across different values for matrix dimensions in the noise_elimination problem. The model was trained on 10 × 10 matrices and was evaluated on larger matrices. Let *solved* instances here refer to those instances that the model finds a conflict-free matrix by flipping at some coordinates. We observed that the model flipped at most 3 coordinates for all solved instances in this experiment. The number of solved instances for each matrix dimension relative to that corresponding to 10 × 10 matrices can be thought of as a measure of generalizability. Noise rates for 10 × 10 matrices are *α* = 4 × 10^−4^ and *β* = 0.02. Noise rates for the rest of the matrices are tuned in a way that we ensure the expected number of flipped entries stays fixed as the dimensions grow.

### 3.5 Application to real data

We also applied our method for no_branching problem to three real datasets for which previous phylogenetic analyses reported highly concordant trees of tumor evolution. The first dataset consists of 16 single-cells from a Triple Negative Breast Cancer (TNBC) patient. Originally it was made available in (Wang et al. 2014b), and here we focused on the analysis of 18 mutations previously used in (Malikic et al. 2019a). The evolutionary history of these mutations was first (indirectly) reported in the original study, and the same results were later obtained by the use of B-SCITE (Malikic et al. 2019a). As shown in (Malikic et al. 2019a), the reported tree has high support from both single-cell and bulk data.

Second, we analyzed an Acute Lymphoblastic Leukemia (ALL) Patient 6 dataset from (Gawad et al. 2014), where 146 single-cells were sequenced and 10 mutations reported. Similar to the TNBC dataset, the evolutionary history of these mutations was thoroughly studied in the original study and later in (Kuipers et al. 2017).

Third, we applied our method to the data from Patient 1, diagnosed with colorectal cancer that metastasized to the liver, published in (Leung et al. 2017) (note that in (Leung et al. 2017) this patient is also labeled as CRC1 or CO5). In total, 16 mutations were detected in the original study, and the evidence for presence of at least one of these mutations was found in 40 single-cells from the primary (colon) and 32 single-cells from the metastatic (liver) site. Detailed analysis of the evolutionary history of this patient was also performed, and a tree of tumor evolution inferred by the use of SCITE (Jahn et al. 2016).

In order to apply our approach for the no_branching problem on these datasets, we trained four separate models with the same architecture described in Section S1.1. The dimensions of the input were set to 16 × 18 for the first dataset and 146 × 10 for the second dataset. We trained two models to apply to the third dataset. The first model was trained with input dimensions set to 40 × 16 to process the submatrix corresponding to the cells from the primary site. The second model was trained with input dimensions set to 72 × 16 to process all cells (from both primary and metastatic sites).

In the training phase, only simulated instances (described in Section S1.1) were used. Note that one of the drawbacks for the model based on a feed-forward neural network without any recurrent layer, described in Section S1.1, is that input dimensions are fixed. One can use padding or copying techniques to adjust the dimensions to any number of dimensions that a pre-trained model requires, as long as the number of dimensions of the input is smaller than that required by the model. However, this may have an adverse impact on the accuracy; training with the same number of dimensions typically leads to better results.

According to our results, the tree corresponding to the first dataset contains at least one branching event (i.e., has non-linear topology), with probability 0.97. This result is highly consistent with previous analyses of this dataset, wherein each analysis, several branching events in the tree of tumor evolution of this patient were reported (Malikic et al. 2019a, 2020, Ramazzotti et al. 2019, Singer et al. 2018, Wang et al. 2014b).

For the ALL patient, the probability of at least one branching event is only 0.21, suggesting a linear topology. This topology was also reported in the original study and later inferred by the use of SCITE (Jahn et al. 2016) in (Kuipers et al. 2017). Small but non-zero branching probability of 0.21 for this patient can be explained by some evidence for a recurrent mutation in gene SUSD2 presented in (Kuipers et al. 2017), which may potentially be due to a local branching event (involving only two mutations at the leaf level).

For the third dataset, the tree reported in the original study (Leung et al. 2017) implies linear evolution at the primary tumor site (i.e., when we restrict our analysis to the cells originating from the primary site). However, when primary and metastatic cells are combined, the metastasis-specific mutations form a separate branch resulting in a tree with branching topology. In order to test whether our method can successfully distinguish the two cases, we run it on two different inputs: the first input consisting only of single-cells sampled from the primary site and the second input consisting of cells from both primary and metastatic sites. Our models′ probabilities of having linear topology reported on the first and the second inputs equal to 0.83 and 0.22, respectively. These values are highly consistent with the results presented in (Leung et al. 2017).

## 4 Discussion

This paper introduces deep learning solutions to the three problems related to tumor phylogeny inference from SCS data, i.e., given a binary genotype matrix obtained from SCS data, with rows representing singlecells and columns representing putative mutations, and false positive and false negative noise rates of the data: (1) Decide whether the most likely tumor phylogeny has a linear topology or contains one or more branching events, (2) Decide whether the matrix satisfies three gametes rule (and thus has a matching perfect phylogeny with genotypes of leaves of the phylogeny matching rows of the matrix), (3) Find the most likely tumor phylogeny based on the observed genotype matrix.

Note that there can be multiple equally likely solutions to the tumor phylogeny reconstruction problem and currently no available combinatorial technique can guarantee the inference of all of these solutions (in fact, all available techniques aim to infer a single most likely tumor phylogeny). Preliminary simulations on small datasets seem to suggest that the most likely tumor phylogeny inference problem possibly has a single optimal solution provided the noise on the input SCS data is independently applied to the entries of the input genotype matrix. However, a larger-scale study needs to be conducted to obtain a better understanding of the solution space.

### Method

Please refer to Section S1 in the supplemental file for detailed descriptions of methods.

### Limitations of Study

Our proposed approaches for no_branching problem and noise_inference problem have the drawback that the input dimensions are fixed. We present a few remedies for this issue in Section 3.5. Furthermore, we did not perform any empirical running time comparison between our approach for noise_inference problem and the linear-time algorithm introduced in (Gusfield 1991), which is already asymptotically optimal. In addition, the real datasets explored in Section 3.5 for the no_branching problem were too large, or their corresponding noise rates were too high for our noise_elimination method, and thus we were not able to include any results. Moreover, our approach does not improve the running time for obtaining a conflict-free matrix compared to the previous methods (e.g., PhISCS (Malikic et al. 2019b)).

## Author Contributions

S.C.S. identified the problems discussed in this work and foresaw the potentials for the use of machine learning methods for solving them. M.H.E. and E.S.A., with the help and supervision of S.C.S, S.M., and R.K., developed and implemented the methods and ran the experiments. S.M. also provided guidance on the real-world datasets choice and design of the related experiments. R.K. advised the group on the development of machine learning methods and experiment design. All authors contributed to the writing of the manuscript.

## Acknowledgements

This work is supported in part by the Intramural Research Program of the National Institutes of Health, National Cancer Institute. This work was also supported in part by Lilly Endowment, Inc., through its support for the Indiana University Pervasive Technology Institute (Stewart et al. 2017). E.S.A. and S.C.S. were supported in part by NSF grant AF-1619081, R.K. was supported in part by NSF grant IIS-1906694, and M.H.E. was supported in part by Indiana U. Grand Challenges Precision Health Initiative.

## Resource Availability

### Lead Contact

Feel free to contact S. Cenk Sahinalp at for further correspondence.

### Materials Availability

There are no physical materials necessary to understand or reproduce the claims of this work.

### Data and Code Availability

The code for the construction of simulated datasets and running the training and evaluation for all of the networks described in this work is available at https://github.com/algo-cancer/PhyloM.

## Footnotes

1 See (Malikic et al. 2020) for a proof. Interestingly, for a given binary matrix *S*, it can also be shown that having only one of the two sets of inequalities (see the definition of the staircase matrix) satisfied is sufficient to conclude that *S* is conflict-free and implies clonal tree with linear topology. However, this observation does not affect any of the results presented in this work.

2 We found one proof of this in (Weber & El-Kebir 2020 (to appear), which was presented at WABI 2020 conference after our work was accepted for publication.

