## Supplemental information for "Tumor Phylogeny Topology Inference via Deep Learning"

### Supplemental Information
### Tumor Phylogeny Topology Inference via Deep Learning

Erfan Sadeqi Azer, Mohammad Haghir Ebrahimabadi, Salem Malikić, Roni Khardon, S. Cenk Sahinalp

#### S1 Transparent Methods

##### S1.1 Solutions to the Branching Inference Problem

**Training Dataset.** For this problem, we first generate a set of conflict-free matrices such that $\mathsf{Sing}(A) = 0$ for each matrix $A$ from this set (i.e., tree implied by $A$ has a branching topology). We generate this set through the procedure explained in Section 2. In order to generate a second complementary set of conflict-free matrices, for each matrix $A$ from the first set, we describe below how a new random matrix $A^L$ can be constructed with the same number of ones, but $\mathsf{Sing}(A^L) = 1$, i.e., $A^L$ satisfies the no-branching property. This construction not only guarantees the union of these two sets have a balanced number of instances for branching and non-branching (linear) topologies, but also eliminates any *artificial* distinguishing features (e.g., based on the number of ones in the matrix, which is expected to be higher for matrices implying topologies with a lower number of branching events).

Briefly, consider an arbitrary matrix $A_{n\times m}$ from the set of matrices satisfying no-branching property generated above. Assume that $A$ contains exactly $l$ entries equal to 1. In order to generate related matrix $A^L$ we first produce a random non-decreasing sequence consisting of $n$ non-negative integers $u_1, u_2, \ldots, u_n$ which add up to $l$ and each is upper-bounded by $m$. For each $i \in \{1, 2, \ldots, n\}$ and $j \in \{1, 2, \ldots, (m - u_i)\}$ we set $A^L[i, j] = 0$. All other entries of $A^L$ are set to 1. The matrix $A^L$ generated using this procedure obviously contains exactly $l$ ones. Furthermore, $A^L$ is also a staircase matrix and therefore it has the no-branching property.

Before adding it to our training dataset, we modify the matrix $A^L$ generated above by randomly permuting its rows and columns (this obviously does not affect the topology implied by $A^L$ nor the number of ones that it contains).

We combined the two sets of matrices and used them for training without adding noise. However, we evaluated the trained model on both noisy and noise-free inputs.

**Network Structure.** A two-layer fully connected artificial neural network is used with either 10 or 100 hidden neurons. The input layer corresponds to an arbitrary-size batch of vectors with length $n \cdot m$. We used $\mathsf{tanh}$ as the activation function. To avoid overfitting, a drop-out (Srivastava et al. 2014) rate of 0.9 was used. Note that the architecture had a moderate level of sensitivity to the hyperparameter settings (i.e., the number of neurons and the drop-out rate). The output of this network is one value indicating the probability of the input matrix having no-branching property. During training, a binary cross entropy loss (Bishop 2006) is used to compare the predictions against binary labels given as the ground truth.

##### S1.2 Solutions to the Noise Inference Problem

In order to solve the noise_inference problem, we formulated it as a classification task through supervised learning.

**Training Dataset.** As described in Section 2, we first generated a set of conflict-free matrices $A$, i.e., $\mathsf{lcf}(A) = 0$, and then for each matrix $A$ we obtained its noisy version $A'$ with predetermined noise parameters

$\alpha$ and $\beta$ so that $\mathsf{lcf}(A') = 1$. The combined set of matrices $A$ and $A'$ forms a balanced binary dataset of noisy and noise-free matrices as required for classification purposes.

For some of our experiments, we preprocessed each input matrix as per (Gusfield 1991) by sorting its columns in non-decreasing order based on the binary number each represents (read from top to bottom). E.g., this would move column $\begin{bmatrix}0\\1\\1\end{bmatrix}$ to the left of column $\begin{bmatrix}1\\0\\0\end{bmatrix}$ as $(011)_2 = 3 < (100)_2 = 4$. Sorting columns was shown to help checking whether a matrix is conflict-free or not in non-ML settings (Gusfield 1991). We show that it also helps ML strategies (see Section 3).

**Network Structure.** For this problem, we use a two-layer fully connected network with 100 neurons in each layer. The input layer corresponds to an arbitrary-size batch of vectors with length $n \cdot m$. The Sigmoid function was used as the activation function for both layers. We used a drop-out rate of 0.2 for each layer for preventing our network from overfitting. The output of this network is one value indicating the probability of the input matrix being conflict-free. During training, a binary cross entropy loss (Bishop 2006) is used to compare the predictions against binary labels given as the ground truth.

#### S1.3 Solutions to the Noise Elimination Problem

This section presents our approach for solving the noise_elimination problem and thus contains our major contributions from a methodological standpoint.

##### S1.3.1 Input Format

Recall that input to this problem is given as a binary matrix $A'$. We transform $A'$ to a new matrix $A''$ as shown below, before feeding it to the neural network. Each row of $A''$ corresponds to exactly one of the entries of $A'$. The first two columns of a given row in $A''$ respectively represent the row and the column of the corresponding entry in $A'$. The third column represents the noise level and depends on the actual value of the entry - which is represented in the last column. In the simplest case, the value of the third column is either $\alpha$ - if the entry has value one, or is $\beta$ - if the entry has value zero. In the more general case, the user can specify a distinct false positive or false negative rate for each entry of the input matrix $A'$. As depicted below, this transformation is key to our ability for training the neural network with matrices of varying shapes/dimensions:

$$\underset{A'_{n\times m}}{\begin{bmatrix} a'_{1,1} & a'_{1,2} & \dots & a'_{1,j} & \dots & a'_{1,m} \\ \vdots & \vdots & \ddots & \vdots & \ddots & \vdots \\ a'_{i,1} & a'_{i,2} & \dots & a'_{i,j} & \dots & a'_{i,m} \\ \vdots & \vdots & \ddots & \vdots & \ddots & \vdots \\ a'_{n,1} & a'_{n,2} & \dots & a'_{n,j} & \dots & a'_{n,m} \end{bmatrix}} \rightarrow \underset{A''_{n\cdot m\times 4}}{\begin{bmatrix} 1 & 1 & f(a'_{1,1}) & a'_{1,1} \\ 1 & 2 & f(a'_{1,2}) & a'_{1,2} \\ \vdots & \vdots & \vdots & \vdots \\ i & j & f(a'_{i,j}) & a'_{i,j} \\ \vdots & \vdots & \vdots & \vdots \\ n & m & f(a'_{n,m}) & a'_{n,m} \end{bmatrix}}$$

where,

$$f(a'_{i,j}) = \begin{cases} \alpha & \text{if } a'_{i,j} = 1 \\ \beta & \text{if } a'_{i,j} = 0. \end{cases} \tag{4}$$

Note that the values of $\alpha$ and $\beta$ are typically estimated as part of single-cell data analysis (e.g., by considering coverage across heterozygous sites) (Gawad et al. 2014, Leung et al. 2017, Wang et al. 2014a).

While the value of $\beta$ in most of the available datasets is within the range from 0.1 to 0.3, the value of $\alpha$ is usually lower than 0.01.

###### S1.3.2 Method

Our approach is similar to the algorithm in (Bello et al. 2017), which follows the model-free policy-based reinforcement learning framework, and was developed to solve TSP and knapsack problems. This framework requires a *cost function* to be defined on the space of potential outputs for the network. In the training phase, the optimization algorithm tries to minimize this cost function. Notice that this will be the only source of supervision we provide to our model for training purposes. Importantly, we do not need to know or calculate through alternative methods the ground truth $A$, which is more expensive to infer in comparison to calculating our cost function as described below.

The cost function we came up with is defined for a given input matrix $A'$ and a potential output matrix $X$, with entries $x_{i,j}$, and has two terms. The first term, $\mathsf{NumberOfConflicts}(X)$ is equal to the number of distinct conflicts present in $X$. As such, it will have value 0 if $X$ is conflict-free:

$$\mathsf{NumberOfConflicts}(X) = \sum_{c_1\in[m],c_2\in[m],c_1<c_2} \Big( \sum_{r_1\in[n]} [(x_{r_1,c_1},x_{r_1,c_2}) = (0,1)] \cdot \sum_{r_2\in[n]} [(x_{r_2,c_1},x_{r_2,c_2}) = (1,0)] \cdot \sum_{r_3\in[n]} [(x_{r_3,c_1},x_{r_3,c_2}) = (1,1)] \Big). \tag{5}$$

In addition to the $\mathsf{NumberOfConflicts}(X)$, we use the negative log-likelihood of $X$, introduced in Equation (2), as the second term of the cost function $\mathsf{Cost}(X) : \{0,1\}^{n\times m} \to \mathbb{R}^+ \cup \{0\}$, which is defined below:

$$\mathsf{Cost}(X) = \mathsf{NumberOfConflicts}(X) - \gamma \cdot \ln(P(A'|A = X, \alpha, \beta)) \tag{6}$$

where $\gamma > 0$ is a hyper-parameter for establishing a trade-off between the two terms.

Recall that the objective of our optimization problem is to compute a matrix with the highest likelihood among all matrices with $\mathsf{NumberOfConflicts}(X) = 0$. Equation 6 achieves this goal when $\gamma$ is set to a small value (e.g., it can be easily shown that any value that is smaller than $-\frac{1}{2n\ln\beta}$ is sufficiently small to be set as a value of $\gamma$).

The core architecture we use here, within the reinforcement learning framework, is a *pointer* network (Bello et al. 2017). Figure S1 shows the components of the pointer network and the flow of information when it is used for evaluation, as summarized below. The additional connections used for training, e.g., for back-propagation, are not shown in the figure.

The embedding layer embeds the input array of shape $(n \cdot m) \times 4$ into an array of shape $(n \cdot m) \times d$, where $d$ is the embedding length used in the network. The encoder consists of $q$ layers of convolution (Fukushima 1980). Each one of the $(n \cdot m)$ rows (with length $d$) of the embedded array goes through the $q$ convolution layers to produce a length $d$ vector placed in the buffer as a new row. In our experiments, the values of $d$ and $q$ were set to 8 and 2, respectively. The decoder is a long short-term memory (LSTM) (Hochreiter & Schmidhuber 1997) layer and the attention layer (Bahdanau et al. 2015) is a network of fixed size. Together they compute the output of the neural network. Specifically, the attention layer iteratively calculates a real valued *score* function on $d$ dimensional vectors. In each iteration, this function is applied to each row of the buffer to obtain its score, which is then used to pick a single row from the buffer (corresponding to a row of the input to be flipped), to be fed to the LSTM. The output of the LSTM is another vector of length $d$, which is used to update its memory and is then fed to the attention layer, to be used to update the score function.

The cost function we described earlier is employed both in updating the LSTM as well as the score function of the attention layer. Note that the score function is not trivially applied only to the rows of the buffer that have not been picked earlier. Those that have been picked are automatically assigned a fixed low score. A soft-max function is then used on these scores to get a probability distribution over the rows of the buffer. In the next iteration, the row to be fed to LSTM is sampled according to this distribution.

This neural architecture is used to decide and evaluate a set of proposed flips through the cost function. More specifically, the buffer row selected by the attention mechanism at each iteration is considered to be the proposed flip location. We combine all the proposed flips to $A'$ to get the output matrix $B$. Then the value of $\mathsf{Cost}(B)$ is calculated and used as feedback for the network. Since the architecture is trained via policy gradients, the total cost function is sufficient as the feedback signal, and the learned policy, implicit in the network and attention mechanism, will attempt to minimize the cost of the output $B$.

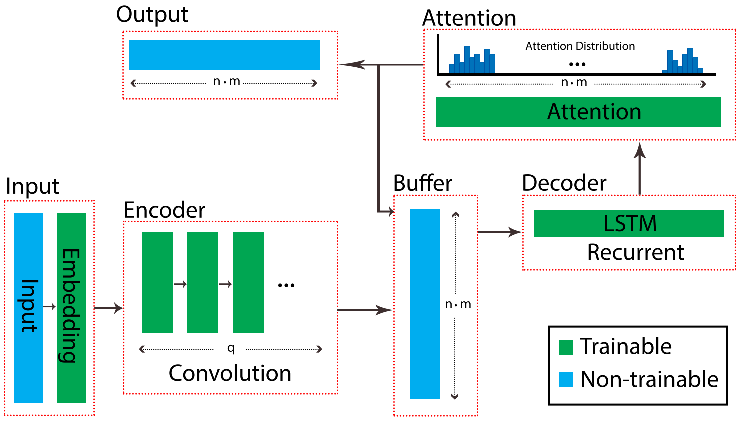


Figure S1: The architecture of the network used for the noise_elimination problem. The roles for the inner-components of the network including embedding layer, encoder, decoder, and attention mechanism interacting with the buffer, are explained in the text. In each training epoch, a batch of input matrices is fed to the network. Each input matrix $A'_{n\times m}$ in a batch, is preprocessed to an $(n\cdot m)\times 4$ matrix as described in Section S1.3.1. The output of the attention layer, right before the final output layer, is a function that forms a distribution over the $n\cdot m$ entries of the input matrix $A'$ (after having been processed through the embedding layer and the encoder). The attention layer iteratively forms the final **output** of the network, which is a set of entries of $A'$ to flip.

#### S2 Experimental setup

All the experiments in this work are performed using Carbonate, a computer cluster at Indiana University. We used deep learning (DL) nodes in Carbonate. These nodes feature 12 GPU-accelerated Lenovo ThinkSystem SD530, each equipped with two Intel Xeon Gold 6126 12-core CPUs, two NVIDIA GPU accelerators (eight with Tesla P100s; four with Tesla V100s), and 192 GB of RAM. A Red Hat Enterprise Linux Server 7.7 is running on these nodes, and we used Python 3.7.3 :: Anaconda, Inc., Tensorflow 1.13.1, and Keras 2.2.4.

#### S3 Details of simulating genotype matrices

In order to generate genotype matrices used as the input, we first run ms simulator (Hudson 2002), which generates conflict-free matrices under the infinite sites model. We run the following command to generate conflict-free matrices:

```
>>> ms nsam nreps -s
```

where,

- nreps denotes the number of independent samples to generate (the number of conflict-free matrices)
- nsam denotes the number of copies of the locus in each sample (the number of cells)
- −s denotes the number of segregating sites (the number of mutations)

Let $A$ denote a conflict-free matrix generated by ms. In order to obtain instances of a noisy matrix $A'$ used in our experiments, we *added noise* to $A$ by *flipping* some of its entries. For each entry which equals 1 in $A$, we decide whether to flip it or not by a simple draw from the Bernoulli distribution with the probability of success equal to the false negative error rate of single-cell sequencing experiment. In case of success we set $A'[i,j] = 0$, otherwise $A'[i,j] = 1$. Entries $A'[i,j]$ for pairs $(i,j)$ for which $A[i,j] = 0$ are obtained analogously by considering false positive error rate of single-cell sequencing experiment as the success probability in the Binomial distribution.

Recall that we also check each matrix $A'$ labeled as a matrix containing a conflict to verify that it has at least one *conflict*. If it does not, the above procedures are repeated until a new matrix $A'$ containing conflict is produced.

#### S4 Discussion of the sensitivity of the experiments to hyperparameters

There are a few parameters in our methods that can take values from their domain sets. We run an exploratory experiment and observe that there is no high sensitivity when these parameters are changed. Here, we provide a brief discussion of three main parameters.

**$\gamma$ in cost function.** We performed some preliminary experiments (not mentioned in Section 3.4) on matrices with size $10 \times 10$ for $\gamma$ in range $[0.004, 4]$ to explore the impact of this parameter on accuracy. We observed that with $\gamma = 0.004$ the accuracy is 86%. Similarly for $\gamma$ with values 0.04, 1, and 4 the accuracy is 83%, 80%, and 84% respectively.

**Drop-out rate.** The neural networks in Sections S1.1 and S1.2 use drop-out technique to avoid overfitting. We tried several values from the range $[0.1, 0.9]$. We choose to use 0.9 within the networks used in Section S1.1, and 0.2 in Section S1.2. For example in the noise_inference problem using our sorted dataset of size $10 \times 10$ with $\alpha = 0.002$ and $\beta = 0.1$, when the activation function was Sigmoid and the model was trained only for 20 epochs, for drop-out rates of 0.2, 0.5, and 0.8 the evaluation accuracies 88%, 85%, and 78% could be achieved respectively.

**Activation Function.** We tried common activation functions, e.g., ReLU, tanh, and Sigmoid, within the neural networks in Sections S1.1 and S1.2. We choose to use tanh within the network in Section S1.1, and Sigmoid in Section S1.2. We observe that these two functions are generally performing better than ReLU. For example in the noise_inference problem using our $10 \times 10$ sorted dataset with $\alpha = 0.002$ and $\beta = 0.1$ we achieved 82% and 80% evaluation accuracy while ReLu and tanh were used as activation functions, respectively. For the dataset with the same matrices but unsorted, the evaluation accuracy was 68% when ReLu is used and 67% when tanh is used as the activation function.

#### S5 Training time

In this section, we report the time it takes for training each of three neural networks that correspond to the three problems that we attempt to solve in this work.

**Branching Inference.** The training time for $2K$ matrices, including $1K$ from each class of size $100 \times 100$ for 200 epochs, was 995.21 seconds on our machines. With the same dataset size and the number of epochs, it took 1547 seconds (around 25 minutes) to train matrices with $500 \times 500$ dimensions.

**Noise Inference.** For this problem training on $2M$ matrices, $1M$ each class, of size $10 \times 10$ for 500 epochs took 65940 seconds. For the same number of training and evaluation matrices of size $25 \times 25$ and the same number of epochs, the running time was 67979 seconds. It is worth mentioning that noise rates do not affect the training time.

**Noise Elimination.** In this problem, the network was trained on batches consisting of 128 unique matrices. On each batch, the network was trained only once. The training time for all 14000 batches (i.e., 1792000 matrices in total) was 52380 seconds.
